# A Process-Based Model with Temperature, Water, and Lab-derived Data Improves Predictions of Daily Mosquito Density

**DOI:** 10.1101/2021.09.08.458905

**Authors:** D.P. Shutt, D.W. Goodsman, Z.J.L. Hemez, J.R. Conrad, C. Xu, D. Osthus, C. Russell, J.M. Hyman, C.A. Manore

## Abstract

While the number of human cases of mosquito-borne diseases has increased in North America in the last decade, accurate modeling of mosquito population density has remained a challenge. Longitudinal mosquito trap data over the many years needed for model calibration is relatively rare. In particular, capturing the relative changes in mosquito abundance across seasons is necessary for predicting the risk of disease spread as it varies from year to year. We developed a process-based mosquito population model that captures life-cycle egg, larva, pupa, adult stages, and diapause for *Culex pipiens* and *Culex restuans* mosquito populations. Mosquito development through these stages is a function of time, temperature, daylight hours, and aquatic habitat availability. The time-dependent parameters are informed by both laboratory studies and mosquito trap data from the Greater Toronto Area. The model incorporates city-wide water-body gauge and precipitation data as a proxy for aquatic habitat. This approach accounts for the nonlinear interaction of temperature and aquatic habitat variability on the mosquito life stages. We demonstrate that the full model predicts the yearly variations in mosquito populations better than a statistical model using the same data sources. This improvement in modeling mosquito abundance can help guide interventions for reducing mosquito abundance in mitigating mosquito-borne diseases like the West Nile virus.

## 1 Introduction

*Culex* mosquitoes are a primary vector for West Nile virus (WNV) in the United States and Canada [1, 2, 3, 4, 5]. First introduced to the United States in 1999 and to Canada in 2001, WNV is a potentially fatal mosquito-borne disease [1, 2, 6]. *Culex pipiens* and *Culex restuans* are known to transmit WNV in North America [7]; therefore, being able to predict their abundance could provide public health professionals with a system to help anticipate and mitigate disease outbreaks.

We create a process-based modeling (PBM) approach for predicting mosquito abundance that incorporates time-dependent data streams for water levels of nearby streams, ponds, canals, and lakes. The model follows the life-cycle of the mosquito population, and the development is influenced by environmental variables, incorporates temperature, daylight hours, and aquatic habitat availability. The model can predict the impact of temperature, precipitation, and water resource management approaches on seasonal mosquito populations. Reducing the mosquito populations will have a direct impact on mosquito-borne disease transmission.

The mosquito life cycle and virus incubation rates are directly related to temperature. Therefore, many mosquito prediction studies have focused on temperature-dependent approaches [8, 9, 10, 11, 12]. While temperature can capture seasonal trends well, studies have concluded that additional factors must be considered to accurately capture the fluctuation in mosquito abundance over the seasonal trend [9, 10]. Recent studies have addressed these concerns by including temperature and precipitation to better capture the year-to-year variation in mosquito abundance [13, 14, 15, 16, 17, 18]. However, studies have alluded to the fact that assessment of the influence of different rainfall regimens on mosquito populations needs further examination as the rainfall linkage to mosquito habitats could depend on factors such as slope, river routing, and availability of potential habitats for mosquitoes. Some have tackled this by using a lag in precipitation measurements to produce more accurate results [13, 19].

Moreover, methods of capturing mosquito population dynamics themselves vary greatly– including statistical, mechanistic (process-based), and hybrid approaches and various combinations of the mosquito life cycle. Ewing et al. examined the effects of temperature on mosquito populations using four delay-differential equations, which represent each stage of the mosquito life cycle [9]. However, this study did not consider the effects of the availability of standing water on aquatic life stage progression. Similarly, Cailly et al. 2012 developed a model using two systems of ordinary differential equations based on the time of year and ten compartments to comprise the four stages of the mosquito life cycle [20]. Other mechanistic models that use a series of ordinary or delay-differential equations were developed by Wang et al., Lou et al., and Gong et al. [11, 21, 22]. Some include diapause explicitly [23, 20, 8], while others model only within-season dynamics. Statistical approaches to model mosquito populations include, but are not limited to, generalized linear models (GLM) [18], site-specific generalized linear mixed models [16], harmonic analysis, and mixed-effect models [15]. The predictors in these studies include temperature, precipitation, elevation, remote-sensing indices, and land use. It is still unclear what the optimal combination of life cycle attributes, data sources, and methods is to best captures changes in the year-to-year abundance of mosquito populations. The complexity of the inter-dependence of the mosquito life cycle with environmental factors has led to this wide variety of models with variation in accuracy and assessment of the usefulness of different data streams. None of these models have been able to capture the high year-to-year variation in abundance of *Culex* mosquito populations with temperature alone.

*Culex* mosquito development is dependent on surrounding temperature [24, 25, 26, 27, 16] as well as the availability of standing water [15, 28] required for the aquatic stages (egg, larva, and pupa) to develop. The weather affects mosquitoes differently during their life cycle stages. The availability of still water has a more significant impact on the development of eggs, larva, and pupa than on adults. We hypothesize that including water gauge measurements in addition to temperature and precipitation will improve our predictions (see Figure 1). We agree with the conclusion in [12] that age structure should be included. Our model controls the progression from one life stage to the next independently to reflect the actual mosquito development and can guide control measures and account for weather conditions that differ from the previously observed input. It has the advantage of tracking every stage of the mosquito life cycle from eggs through host-seeking females, which is important in determining how many adult females are active in a given period for pathogen spread. Our new model predicts mosquito population abundance based on environmental inputs and mosquito biology alone and does not require new initial conditions every year as many models do, instead explicitly modeling diapause triggered by daylight hours and weather inputs to determine emergence and densities early in the year for our test years.

**Figure 1:**
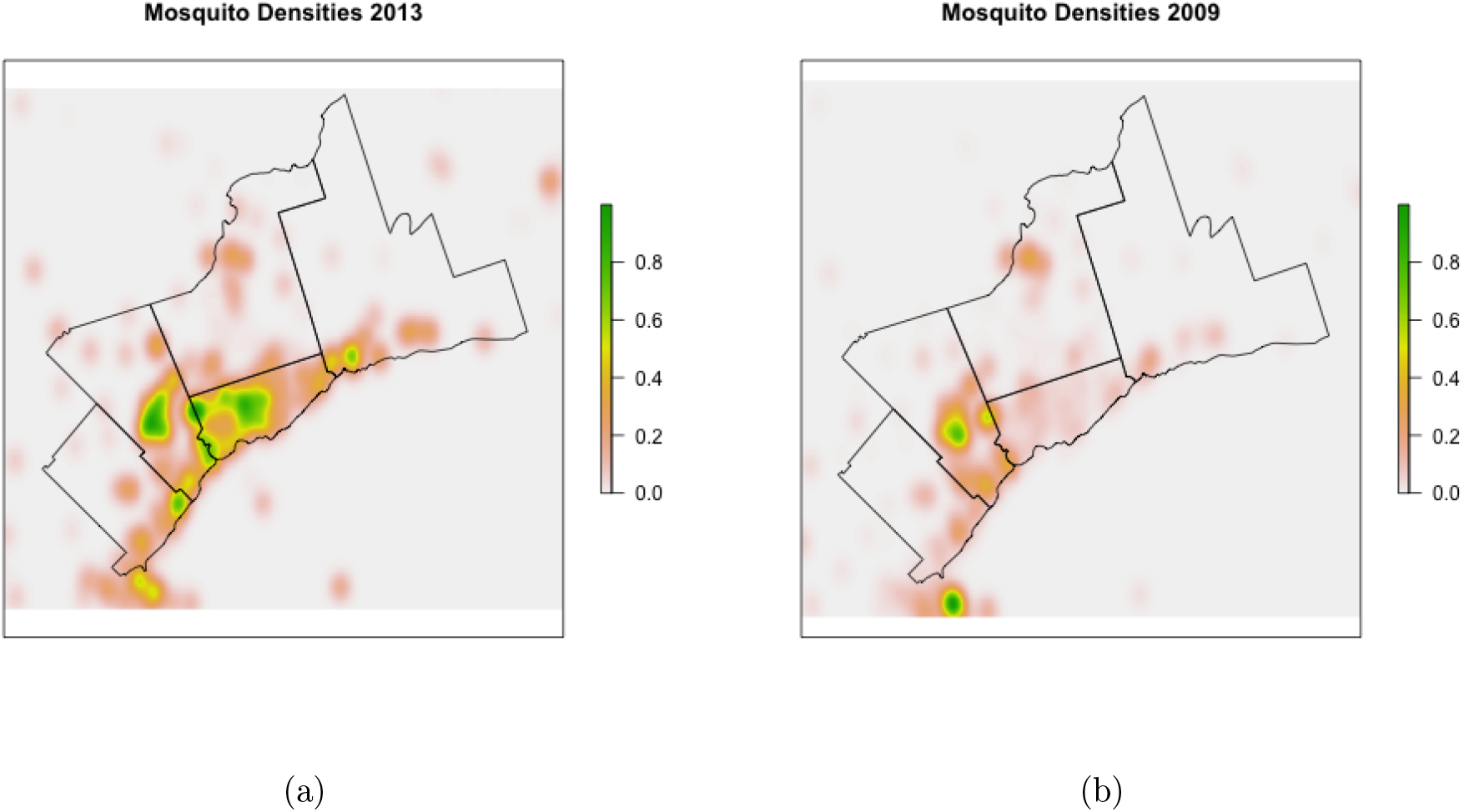
Figure 1(a) is a kernel density estimate of mosquito density in a flood year (2013). Figure 1(b) is a kernel density estimate of mosquito density in a non-flood year (2009)). Flooding often results in higher mosquito trap counts in a wider range of spatial locations in the area, providing motivation for using water levels in addition to temperature for capturing year-to-year changes in mosquito density.

### 1.1 Greater Toronto Area

The most recent studies examining mosquito abundance within the Greater Toronto Area have focused on the Peel Region [18, 15, 8]. Wang et al., 2011 analyzed the association between a gamma-distribution model of mosquito populations with temperature and precipitation in a generalized linear model. This study found temperature to be the most significant factor, while extended periods of precipitation had the greatest effect. They concluded that dynamical equations should investigate other meteorological factors plus all phases of the mosquito life cycle to capture interactive effects between environment and mosquito abundance [18].

Yoo et al., 2016 combined a mixed-effects model with a harmonic analysis of temperature and precipitation to examine association to land use, population density, elevation, spatial patterns, and mosquito abundance. Again this study identified temperature and accumulation of precipitation five weeks prior were the variables of greatest influence [15]. Elevation, the proportion of open space, vegetation, and urban areas had negative correlations with *Culex pipiens* abundance, while NDVI (normalized difference vegetation index) showed only a weak correlation. The authors concluded that this approach fails to capture dynamic interactions between the mosquito life cycle and environmental variables over time. Recently, Yu et al., 2018 exploited a temperature-dependent response function for aquatic and adult life stages over a single season [8]. Their model used temperature alone to predict mosquito life cycle, and the authors concluded that “additional variables needed to be considered to account for the year to year variability in weather”.

Previous mosquito prediction models, developed for the Greater Toronto Area (GTA) and its subsidiaries, focused on precipitation as a proxy for water habitat availability [29, 30, 15]. Studies have concluded that a lag in precipitation, or some other regime of precipitation measurements, better inform mosquito abundance model predictions, [13, 31]. Standing water does not necessarily correlate linearly with precipitation since the amount of flooding caused by a given rainfall volume depends on city infrastructure [32], terrain conditions, and watershed characteristics [33]. Initial analysis of the data (Figure 2 indicated that water station level measurements might be a better signal to capture the seasonal fluctuation of mosquito abundance in Toronto than precipitation measurements.

**Figure 2:**
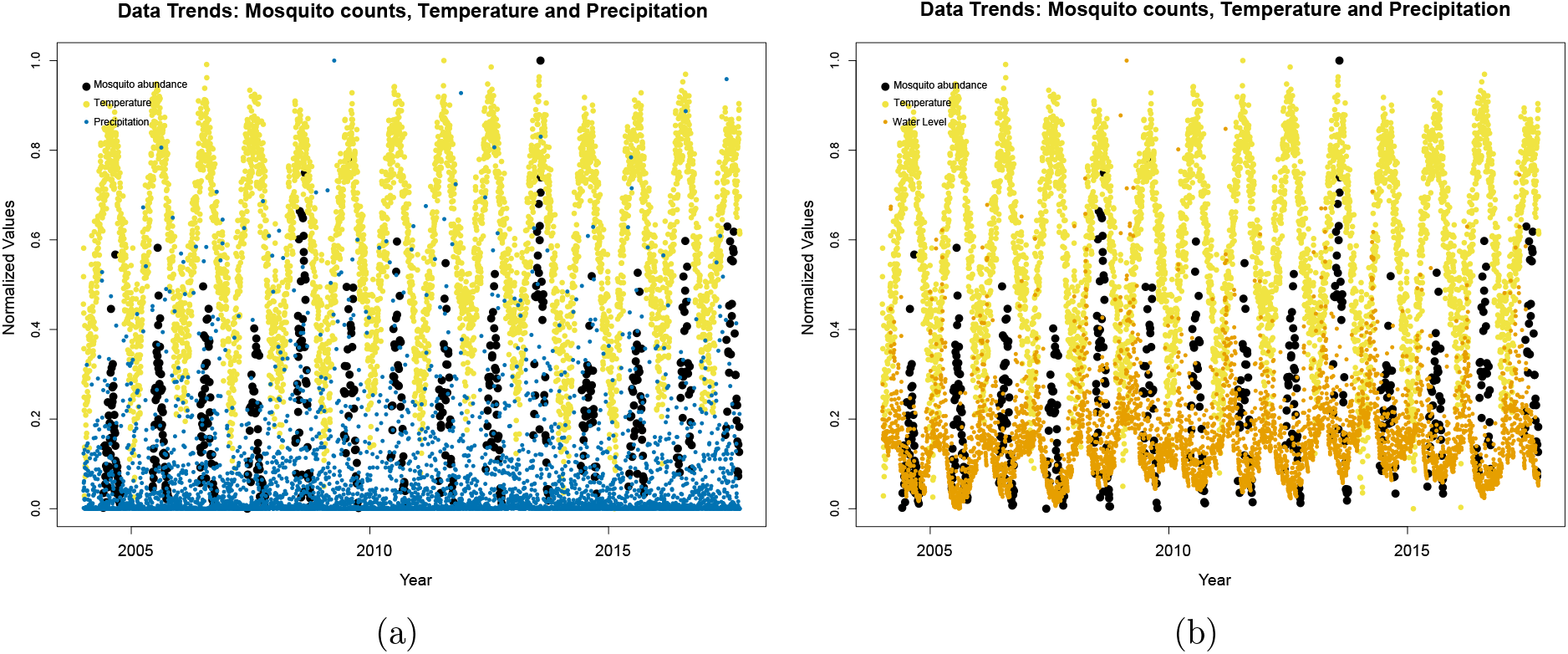
In Figure 2(a) observed mosquito abundance (black), temperature data (yellow) and precipitation measurements (blue) are shown. In Figure 2(b) observed mosquito abundance (black) temperature data (yellow) and water station measurements (orange).

For this study, we used daily municipal water station measurements, daylight hours, and temperature to inform our new process-based model derived from experimental laboratory parameters describing the life stage progression of mosquitoes. This approach extends previous models by combining water gauge levels with laboratory and field data in a mechanistic model. We incorporated water-level and flooding data from the GTA as a proxy to lag measure of precipitation, to predict *Culex pipiens/restuans* mosquito populations (Figure 3). Our PBM is transferable to other locales and mosquito species with appropriate parameterization. To investigate the accuracy of the process-based model, we ran a linear regression model with Gaussian errors to a log-transformed response to compare a statistical model with our process-based model and underscore the need to include the mosquito developmental process. We also fit a.

**Figure 3:**
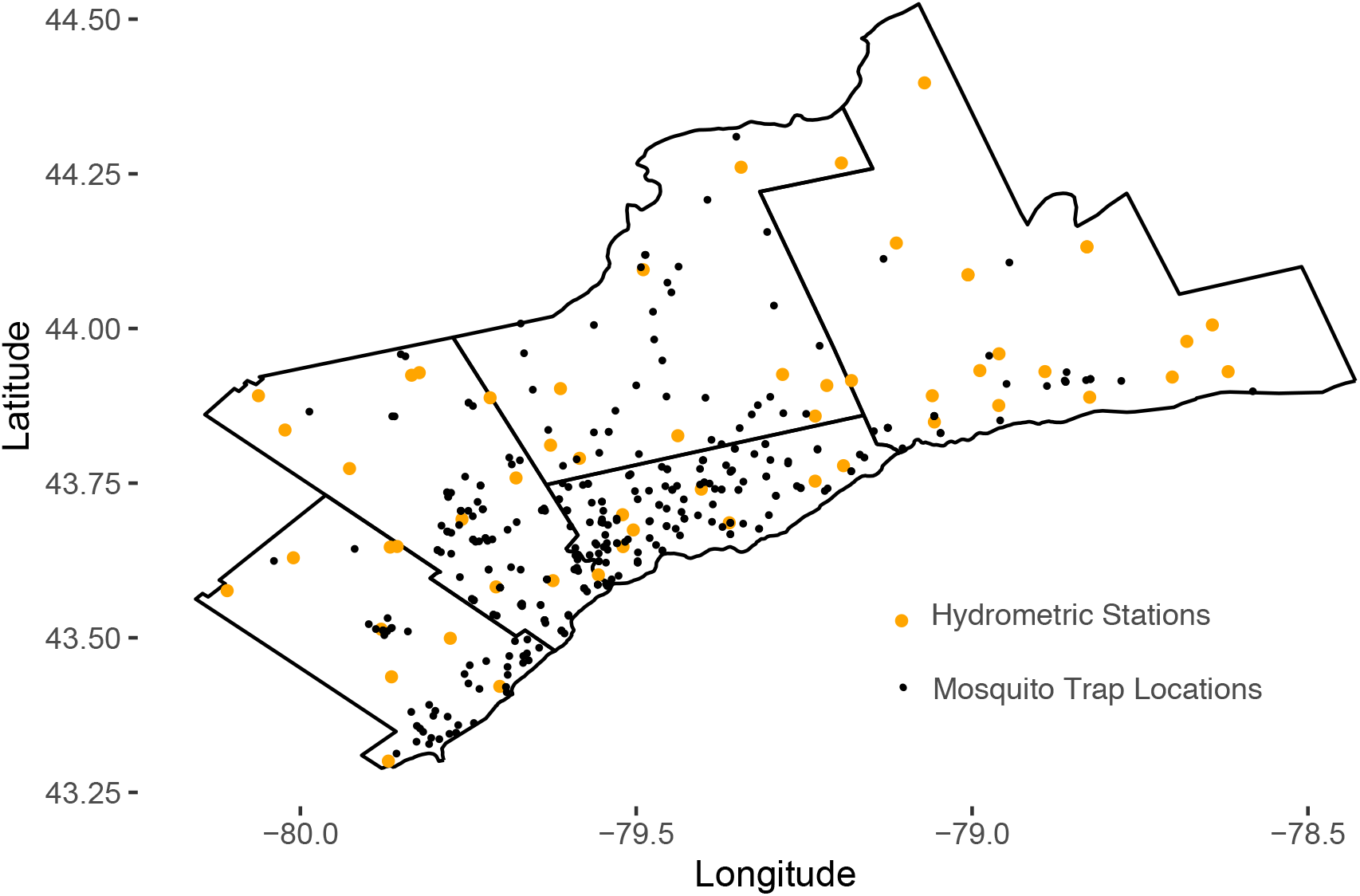
Mosquito trap locations shown on a map of Greater Toronto Area Map (black). Hydrometric stations used for stage gauge water level data are shown in orange.

## 2 Methods

### 2.1 Modeling Approach

We model mosquito abundance in the Greater Toronto Area using two approaches: a process-based model (PBM) and a statistical model. The PBM considers the dependencies of mosquito abundance on exogenous variables through mechanistic equations and tracks the number of mosquitoes in each life stage through time. The statistical model is based exclusively on the correlation between mosquito trap data and the environmental data feeds.

### 2.2 The Process-based Model

Research within laboratory settings has informed understanding of mosquito development and how it depends on temperature and environmental factors (see Figure 4). While these are performed in controlled rather than natural settings, we use the mathematical relationships determined by lab studies within the PBM and adjust for the time-varying environmental inputs in the wild. The PBM incorporates different development rates and death rates for eggs, larvae, pupae, and adult mosquitoes. We describe in the subsequent subsections the algorithm for calculating development progression in and out of the life stages. The final output of the algorithm is a daily prediction for the abundance of the average number of host-seeking female mosquitoes found in a single trap. It produces a continuous time series of the fluctuation in mosquito populations across all 13 years for which environmental data is available. We then compare the model abundance predictions and the observed measurements of average mosquito abundance from mosquito traps (Figure 6). The model is inspired by a partial differential equation approach where mosquitoes develop through life stages in both time and environmental variables via lab-informed kinetics equations.

**Figure 4:**
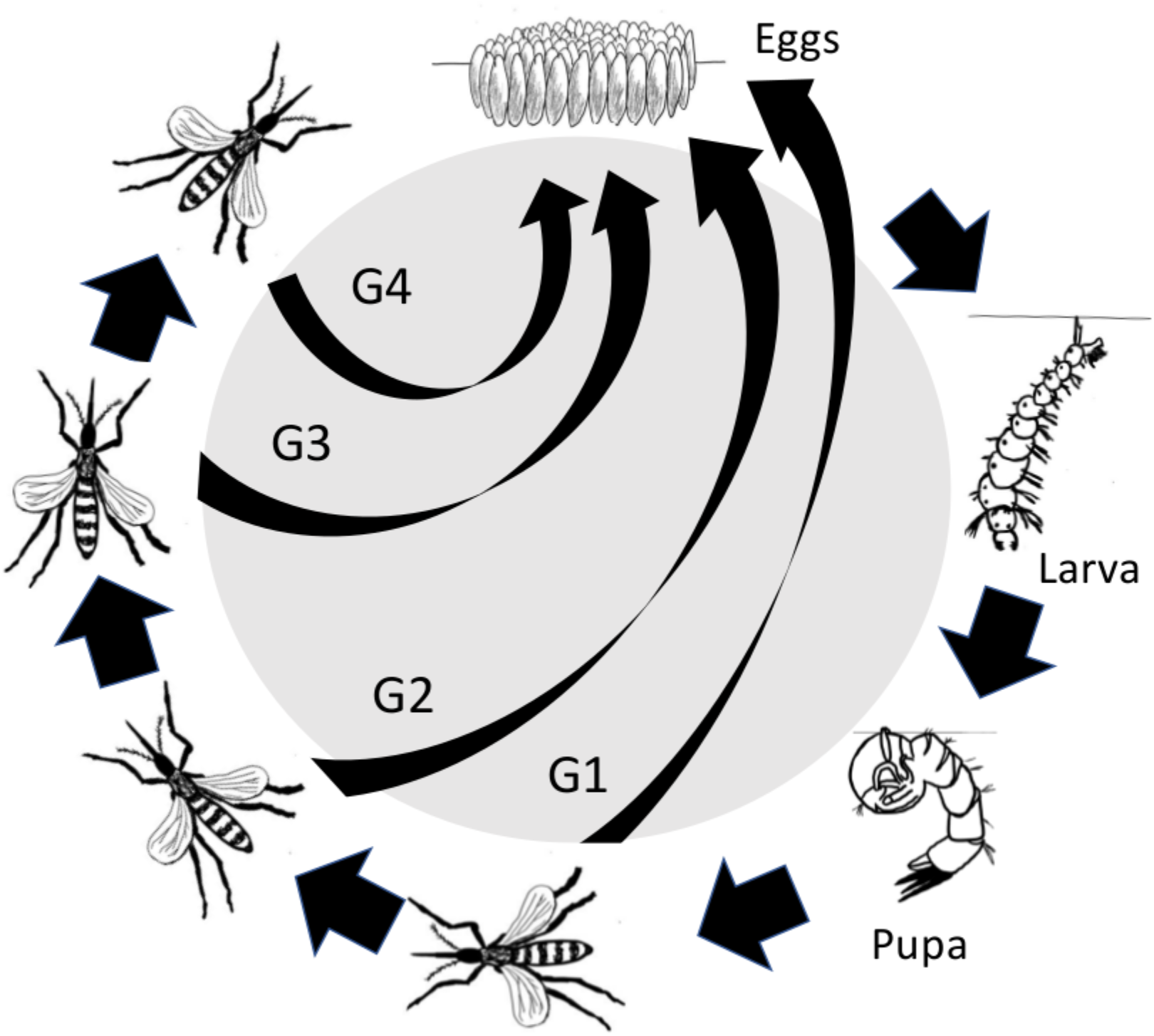
*Culex* mosquito life stages included in our process-based model are eggs, larva, pupa and four adult stages. Over-winter diapause of adults triggered by shorter daylight hours is included, as is competition in the aquatic larva/pupa stage.

**Figure 5:**
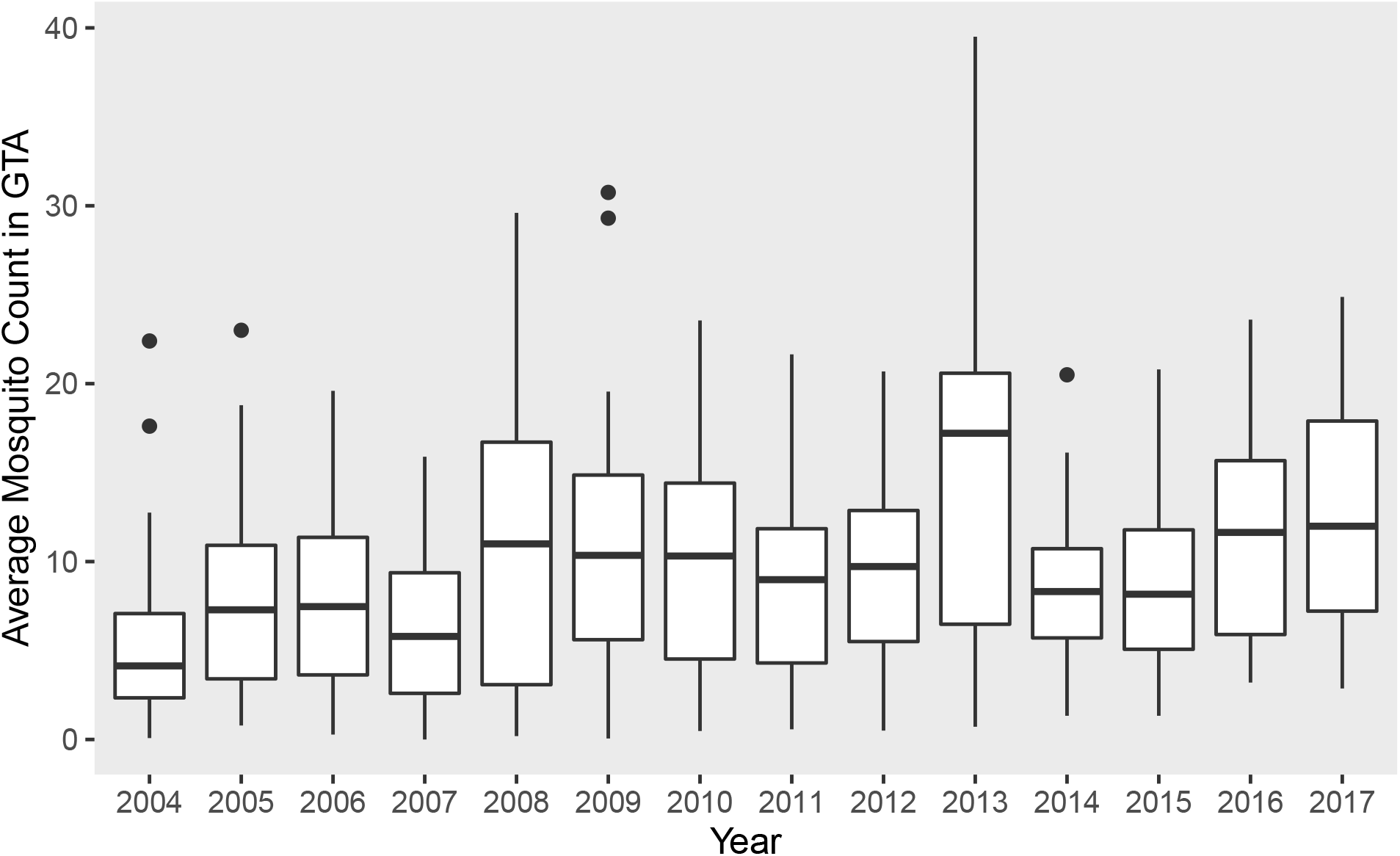
A box and whiskers plot (identifying the 1^st^ and 3^rd^ quartiles, median and outliers) of the arithmetic mean of mosquito counts per trap per year in the Greater Toronto Area.

**Figure 6:**
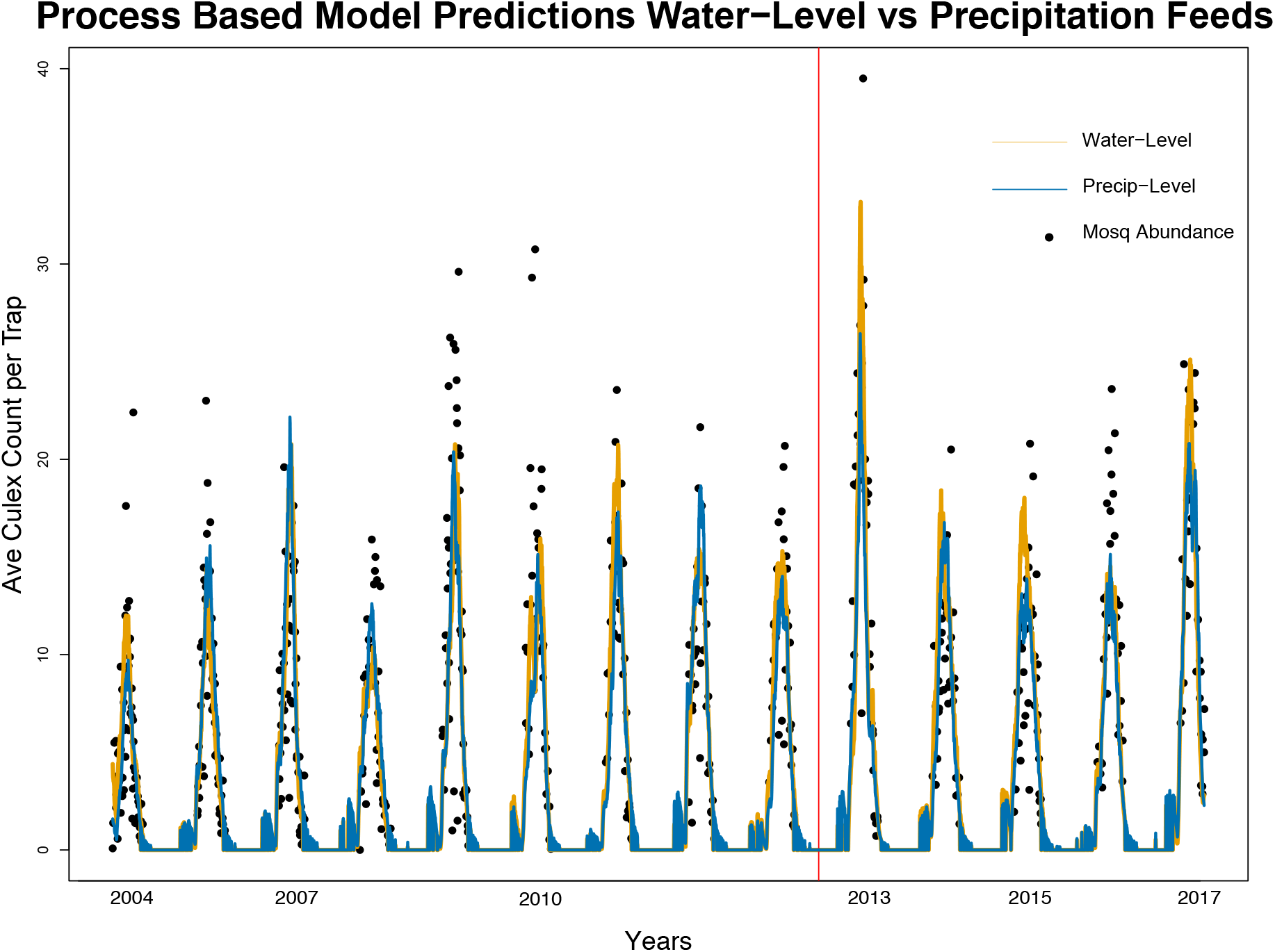
Mosquito abundance predictions from our process-based model using water stage gauge (orange) and precipitation (blue) data overlaid on observed mosquito trap averages (black dots) in the Greater Toronto Area from 2004 through 2017. The vertical red line in both figures indicates the separation between the training data used to fit parameters and the withheld testing data.

#### 2.2.1 Egg Development

We use the Eyring equation [34] to model the developmental progression of mosquito eggs to larva. We assume that eggs do not compete for nutrients but that the development rate, *vE*(*t*), is based on the ambient temperature and time. Thus we use the daily average temperature as the input value for the Eyring Equation:

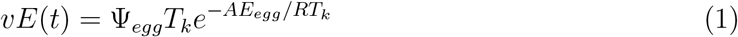

where Ψ*_egg_* and -*AE_egg_* are fit using non-linear least squares of the Erying Equation from laboratory and field studies [35, 25, 36, 22, 37]. *T_k_* is converted from Celsius to Kelvin to be used as an input for the Eyring equation. *R* is the ideal gas constant of proportionality that relates the energy scale in physics to the temperature scale. We impose the restriction that if the daily temperature feed is below 9.154 degrees Celsius, then *vE*(*t*) = 0 [25, 35].

The rate of increase in the egg population depends on the number of adult female mosquitoes calculated to be transitioning from one gonotrophic cycle to the next (and not in diapause). The process for which adult female mosquitoes transition from one gonotrophic cycle to the next will be explained in detail in the adult development subsection.

#### 2.2.2 Larva and Pupa Development

We combine the larva and pupa stages into one group and model the rate of development through this stage via the Briere equation [38]. Again we use identified laboratory behaviors in larva and pupa development and adjust them to depend on environmental real-time data in addition to time:

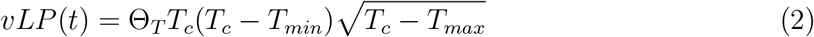

where Θ_T_ is fixed parameter based on laboratory and field data [35, 25, 36, 22, 37] fit to the Briere Equation. *T_min_* and *T_max_* are set as the lower and upper bounds on temperature for which larva and pupa can develop. The daily temperature data feed, *T_c_*, is in degrees Celsius.

We diverge from our previous assumption of neglecting competition to include a density-dependent death rate for the larva/pupa stage, modeled using a quadratic loss differential equation, 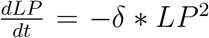, where *LP* is the current size of the larva-pupa population and delta is calculated as

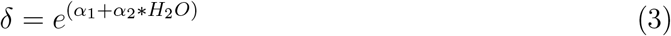

The parameters *α*_1_ and *α*_2_ are fitted to our data. Note that the rate of change, *δ*, used in the quadratic loss function depends on aquatic habitat availability *H_2_O*, incorporating either precipitation or water stage gauge levels as a proxy. Here, we demonstrate that normalized water levels will yield better results in capturing peaks than using normalized precipitation levels. This observation underscores the fact that it is in standing water that *Culex* larva and pupa thrive.

#### 2.2.3 Adult Development

The adult mosquito life stages are identified by the number of times the average mosquito goes through a gonotrophic cycle. At the end of each gonotrophic cycle, the female mosquitoes seek to lay eggs at or near water. Thus the number of newly laid eggs depends upon the number of female mosquitoes ovipositing at any given time. See the Egg Development Section for further details. We will first focus on the method for which the developmental rate of adult mosquitoes, *vAD*(*t*), is calculated. Temperature plays a significant role in development rates for all life stages, including adults. Adult mosquitoes may not transition linearly through all four gonotrophic cycles. There is significant variation in wild adult mosquito lifespan, so we incorporate this variability through a random variable, *M*. We follow closely the method of age distributions used to inform age progression as described in [39]. We diverge slightly from Goodsman et al. in that a Gamma distribution is used for the rate of development of the adult mosquito population, i.e., *M* ~ Γ(*vAD*(*t*), 1). The gamma distribution, Γ(*α,β*) has an expected value of 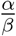. Thus the expected value of the random variable used to model the rate of development *M* is *vAD*(*t*). It is the calculation of the value of *vAD*(*t*) which exploits the rate dependence upon temperature,

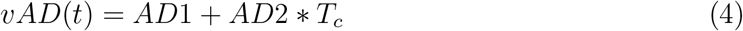

where *AD*1 and *AD*2 are fixed from data, and *T_c_* is the data feed of the daily maximum temperature in degrees Celsius. This will adjust the shape of the gamma distribution based on temperature while at the same time maintaining the observed behavior that extremely high or low temperatures yield slower development rates or may result in early death.

Adult mosquitoes are assumed to have an environmentally dependent death rate. We model the death rate using an exponential decay differential equation where the rate of change within this equation is the Eyring equation [40]. For the adult application of the Eyring Equation, we use the following form:

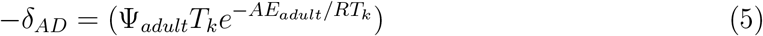

where Ψ*_adult_* and –*AE_adult_* are fit using non-linear least squares of the Erying Equation from laboratory and field studies [35, 25, 36, 22, 37]. *R* is the ideal gas constant. *T_k_* is the temperature data feed per day converted from Celsius to Kelvin.

Diapause is modeled through a logistic regression, which relates the probability of a mosquito being in diapause to the number of daylight hours. The following formula yields the proportion of adult mosquitoes in a given time step which are now in diapause:

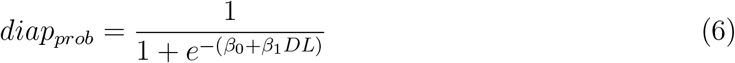

where *β*_0_ and *β*_1_ are fitted from our data, and *DL* is the number of daylight hours per day-step as recorded in the data feed for the Greater Toronto Area.

Eggs are added to the first age of the egg domain at each time step by adult oviposition using the following formula:

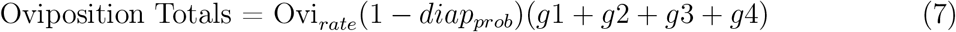

where Ovi_*rate*_ is fitted to our data and *g_i_* is the total number of adult female mosquitoes transitioning from adult gonotrophic stage *i* to *i* + 1 for *i* = 1,…, 4. The *diap_prob_* calculation thus forces the Oviposition Totals to be informed by the maximum average number of daylight hours per day.

The mosquito abundance prediction per day is the sum of the total calculated active mosquitoes. We define active mosquitoes as female mosquitoes which have transitioned from one gonotrophic cycle to the next. These active mosquitoes are actively looking for an oviposition location, e.g. (*g*_1_ + *g*_2_ + *g*_3_ + *g*_4_) for each day.

For detailed tracking of the total number of mosquitoes on any given day, each stage (eggs, larva-pupa, and four adult female stages) is split into 100 evenly spaced compartments through which individuals move based on development rates as described above. Individuals at the end of the compartments will be moved up to the next stage when development rates push them past the end of their current stage.

### 2.3 The Statistical Model

In light of the importance of environmental variables in calculating mosquito abundance, we investigated these data streams without a mechanistic model to see if the data is sufficient to determine the fluctuations in the abundance of mosquitoes. Using a linear model, we aimed to describe the relationships between the mosquito trap data and our environmental variables (temperature, daylight hours, and precipitation/water levels). This method required setting all observed mosquito averages of zero to some small value *ϵ* > 0 as the analysis was performed on a log scale. See Figure 7 for a visualization of the predictive results.

**Figure 7:**
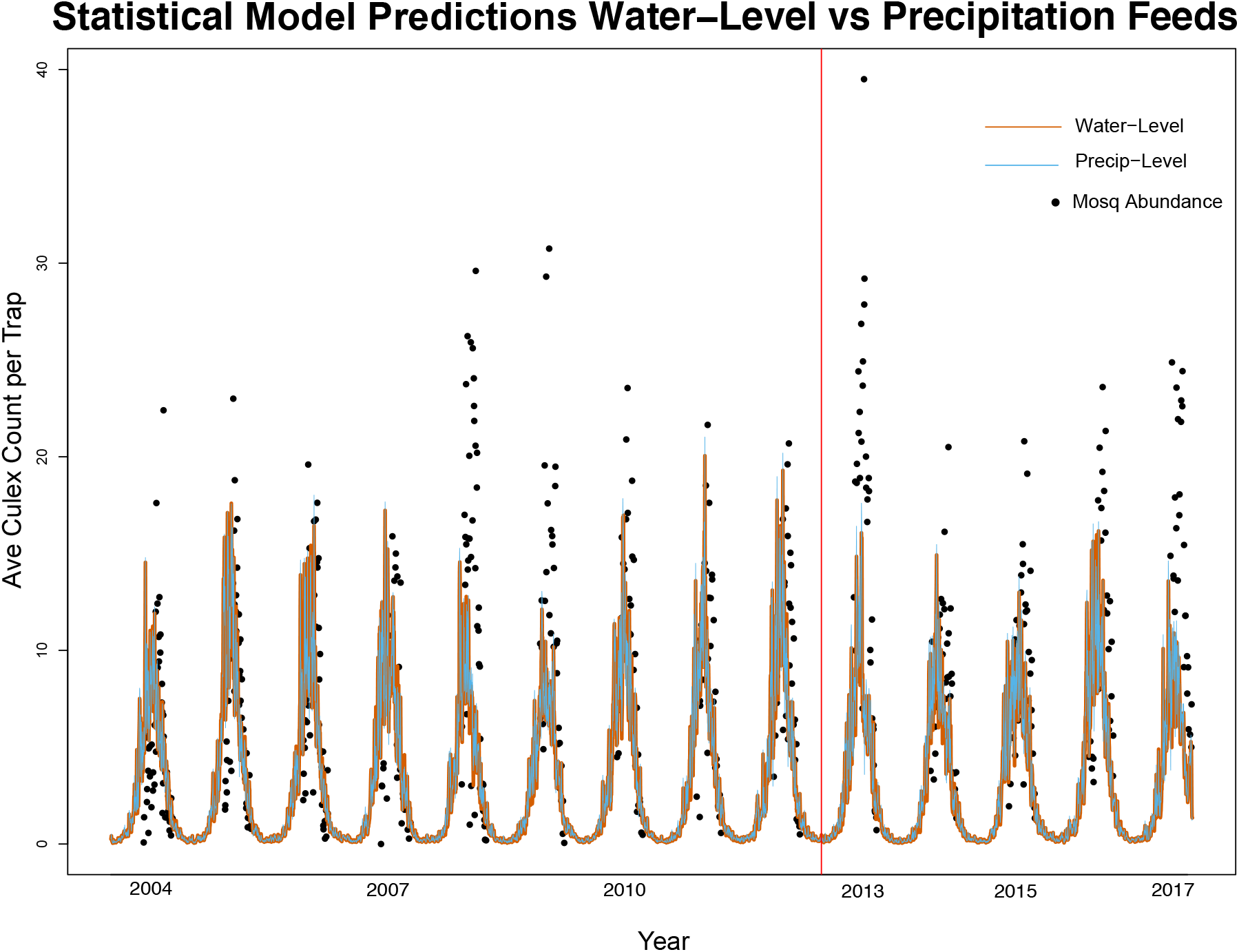
Mosquito abundance predictions from a Generalized Linear Model using water stage gauge (orange) and precipitation (blue) data overlaid on observed mosquito trap averages (black dots) in the Greater Toronto Area from 2004 through 2017. The vertical red line (y = 3300 day-steps equivalently January of 2013) indicates the separation between the training data used to fit parameters and the withheld testing data.

### 2.4 Error Analysis

We computed the root mean squared errors, the mean absolute errors, the correlation coefficients, and the differences between predicted and observed peak number of mosquitoes and the timing of the peak. The PBM consistently outperformed the statistical model, while the water PBM outperformed precipitation PBM for capturing the peak abundance (Table 5) and performed comparably to the precipitation model for daily predictions (Table 1). For the other metrics, they performed similarly.

**Table 1:**
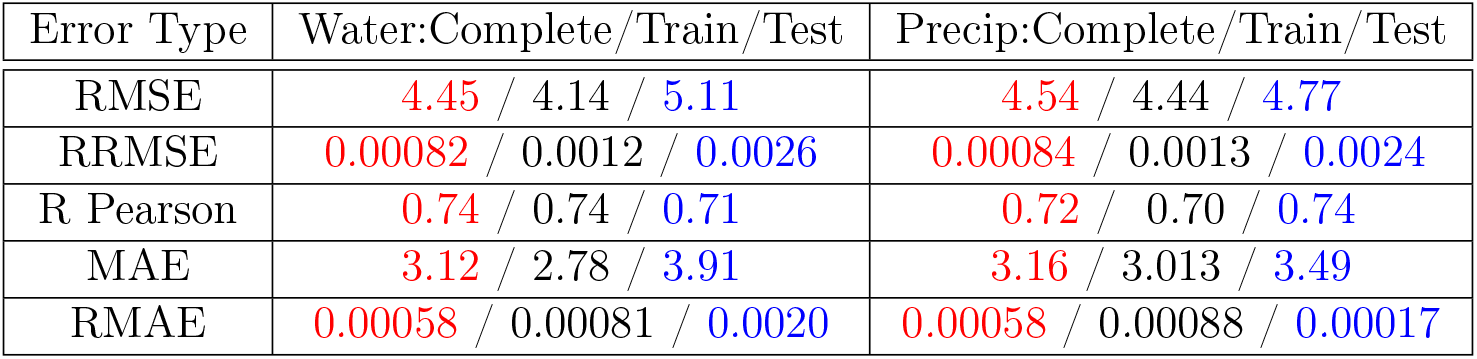
Error Table for the process based model. Red indicates error calculations on the combined training and test data set. Black is the error on the training data set only. Blue is the test data set only.

**Table 2:** Error Table for the statistical model. Red indicates error calculations on the combined training and test data set. Black is the error on the training data set only. Blue is the test data set only.

| Error Type | Water:Complete/Train/Test | Precip:Complete/Train/Test |
| --- | --- | --- |
| RMSE | 6.57 / 5.98 / 7.74 | 6.53 / 5.99 / 7.60 |
| RRMSE | 0.0012 / 0.0017 / 0.0039 | 0.0012 / 0.0017 / 0.0038 |
| R Pearson | 0.37 / 0.38 / 0.39 | 0.37 / 0.38 / 0.38 |
| MAE | 4.60 / 4.14 / 5.63 | 4.57 / 4.17 / 5.48 |
| RMAE | 0.00085 / 0.0012 / 0.0028 | 0.00085 / 0.0012 / 0.0027 |

### 2.5 Parameters

We use average daily temperature, seasonal variation of water station levels, precipitation, and the maximum number of daylight hours to inform the model and fit parameters not found in the literature to mosquito abundance data. To fit the model parameters to data, we employ a nonlinear least squares approach using the Nelder-Mead minimization algorithm to find the parameter estimates that minimize the sum of squared differences between the modeled population trajectory and the observed data. The parameters used in our model are in Tables 3 and 4.

**Table 3:** Description of parameters within the PBM model which were derived from published laboratory findings. **REFS from Devin–Lab Studies**

| Variable | Description | Value |
| --- | --- | --- |
| $\Psi_e$ | Eyring equation for egg development rate | 32270 |
| $AE_e$ | Eyring equation for egg development (neg.) | 41281 |
| $R$ | Ideal Gas Constant | 8.3144 |
| $T_k, T_c$ | data feed of average daily temperatures | K or C° |
| $\Theta_T$ | Briere eqn for larval-pupal development rate | 9.217e-05 |
| $T_{min}$ | Min temp larval-pupal development rate | 9.154 C° |
| $T_{max}$ | Max temp larval-pupal development rate | 36.09 C° |
| $AD_1$ | Adult development rate | -0.111252 |
| $AD_2$ | Adult development rate | 0.013427 |
| $\Psi_{ad}$ | Adult mortality rate | 1.568e+12 |
| $AE_{ad}$ | Adult mortality rate (negative) | 91730 |
| $DL$ | Max number of daylight hours | hrs |
| $H_2O$ | Normalized average water level | unit less |

**Table 4:** Description of parameters within the PBM model which were fitted using the Nelder-Mead Algorithm either using the Water Station Level Data Feed, *H*_2_0 or using the Precipitation Data Feed, Precip.

| Variable | Description | $H_2O$ | Precip |
| --- | --- | --- | --- |
| $\beta_0$ | Diapause intercept | -1.925 | 4.398 |
| $\beta_1$ | Slope proportion in diapause (neg.) | 0.865 | 0.479 |
| $Ovi_{rate}$ | Oviposition rate | 4.729 | 164.964 |
| $Init$ | Initial condition in each age stage | 8.412 | 234.865 |
| $\alpha_1$ | Density-dependent mortality rate | -6.083 | -10.869 |
| $\alpha_2$ | Density-dependent mortality rate | -25.927 | -1548.152 |

### 2.6 Data Sources

All data used for this study is available through the following: Mosquito data for the Greater Toronto Area were obtained from Public Health Ontario’s WNV mosquito database, where public health units trap female mosquitoes every week. A total of 115338 records were collected over 16 years between 2002 and 2017 from 2722 trap sites. In the study region, the mosquito season lasts about 17 weeks between late May and October (weeks 24-40). Day-light hours were calculated based on the day of the year and earth rotation for a single year and duplicated for each year after that. Temperature data was collected using the Rpackage “rclimateca” [41], which fetches data from the Environment Canada climate archives. Information about the boundaries and definitions of the Ontario watersheds were gathered from Land Information Ontario website, last revised on 2010-04-01. Water levels for the riverways and lakes of the Greater Toronto Area were collected from the Rpackage “tidyhydat” [42], which accesses historical and real-time national ‘hydrometric’ stage gauge data from Water Survey of Canada data sources.

While there is variability in the methods used to model mosquito populations, our observations from a detailed literature review show that the data used to inform those models is processed similarly across all methods. Temperature is used as the dominant climatic variable, and precipitation appears almost as frequently. When climate data is collected from multiple stations within the region of interest, daily averages are often used for the model. Most studies use only these variables to construct their models, but others go further to include non-climatic factors such as diapause or varying forms of density-dependent competition [43, 21, 17]. Additionally, the majority of studies use mosquito trap data to calibrate and validate their models. Trap data and climate data for each model always come from the same location. Since our literature review focused on studies which examined *Culex pipiens* and *Culex restuans,* most of the trap data came from temperate regions such as Japan [43], regions of Argentina [13, 12], and the Greater Toronto Area in Ontario, Canada [8, 15, 16, 18]. In most studies, the trap data were collected weekly throughout a period ranging from 1 to 13 years. Similar to the climate data, mosquito counts are averaged when data is collected from multiple traps. When trap data is used to evaluate the predictions made by a model, accuracy is most frequently measured using root mean square error [8], or correlation coefficients [44].

Our mosquito trap observation data includes not only mosquitoes commonly known for carrying WNV such as *Culex pipiens* and *Culex restuans* but also mosquitoes from other genera such as *Aedes* and *Ochlerotatus*. We only considered counts of *Culex pipiens* and *Culex restuans* for our model to keep costs low and to speed up identification. The identifiers grouped both species as a single entry, Cx. pipiens/restuans. Similarly, we filter the data set further to only include those trap sites with lat/long coordinates within the bounds of the Greater Toronto Area. We assume that mosquitoes counted in trap data are active female mosquitoes, as most traps ( 85%) used in our data set are LT traps.

The dates for the observation data start on Jun 6, 2004 and end on Sept 27,2017. We computed the average number of *Culex pipiens/restuans* mosquitoes per trap per day-step, where a day-step = 1 day. The average number of mosquitoes was calculated as the sum of the total number of mosquito counts recorded on a particular day and then divided by the number of traps sites that had observations for that day.

## 3 Results

We compare our PBM predictions with the observed mosquito trap data for the GTA and with a linear statistical model for the same data. We also compare PBM predictions when using stage gauge (water level) data versus precipitation data as proxies for aquatic habitat availability.

In Figure 6, mosquito abundance predictions from our model using daily average temperature, daily maximum daylight hours, and daily normalized averaged water station levels (orange prediction line) is shown with the observed averaged mosquito trap data (black data points). The vertical red line indicates the separation of years for which the data was used to fit our free parameters. The data to the left of the red vertical line was used to estimate the parameters (Table 4). The data to the right of the red vertical line shows the fitted model and mosquito abundance. Figure 6 also displays model predictions using daily average temperature and daily maximum daylight hours now re-fit and predicted with the normalized average precipitation data (blue line).

Our model requires only requires initial conditions for the first year. The remaining 13 years were predicted without defining a new initial condition each season. Most comparable models must re-initialize the mosquito populations every year and require more parameters to fit the data.

The root mean squared error (RMSE) of the water-level PBM is lower than that of the precipitation-based PBM prediction see Table 1. The model RMSE of both types of data are similar as rainfall is correlated with the water levels. Even so, the predicted abundance in Figure 6 indicates the water station level data feed to have a demonstrable effect in capturing the magnitude of the observed mosquito abundance data, especially in flooding years, e.g., 2013 and 2017 (Fig. 6). Using water levels to inform the model beat out using precipitation data in predicting peak levels of mosquito abundance for all years except 2016, see Table 5. The difference between these predictors is less evident in the timing of the peak occurrence.

**Table 5:**
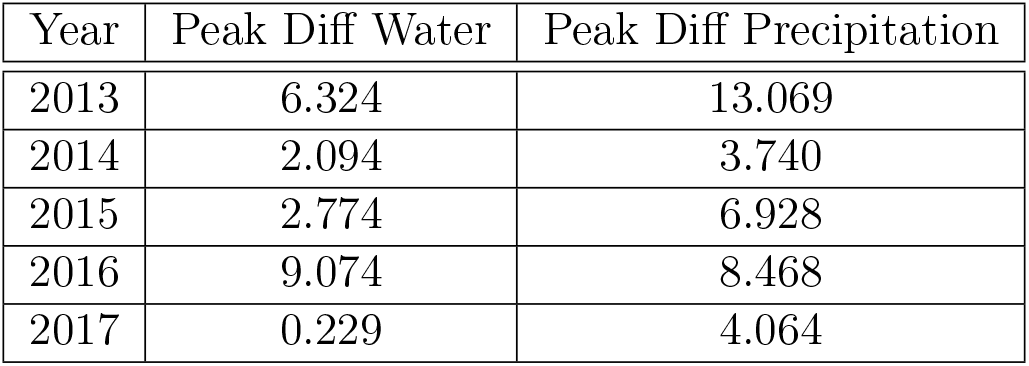
The difference between observed peak mosquito average and the predicted peak values of the corresponding season based on the use of the water data feed versus the precipitation data feed in the Process Based Model are listed as absolute values for the test data years. The model with water gauge data out-performs the model with precipitation data 4 out of 5 years.

We also examined the differences in prediction between including mechanistic effects of exogenous variables on mosquito development versus fitting mosquito populations to the data alone. We fit a linear statistical model to the GTA mosquito trap data using the temperature, daylight hours, and the two aquatic habitat proxies.

We created two linear models to compare the results of using precipitation data and then water station level measurements as proxies for water habitat availability. The two linear models find a correlation between the mosquito population observations (i.e., trap counts) and environmental factors but do not account for causal interdependencies. The results for both linear models are very similar (Figure 7). Predictions from both linear models predictions show the characteristic seasonal behavior of mosquito populations, but neither accurately capture the magnitude of the population for different years.

Both models fail to predict the increase in mosquito population for 2013 and 2017, years with significant rainfall and flooding. The RMSE of the water-level linear model was calculated to be 7.743, and the precipitation-level linear model to be a RMSE of 7.603 (Table 2). Both models were less accurate than the PBM predictions. This comparison supports the observation that a process-based model capturing mosquito population dynamics provides predictive value above and beyond an exclusively data-driven approach.

## 4 Discussion and Conclusion

Though temperature and precipitation levels, when used as drivers or predictors in mosquito population models, provide an excellent correlation to observed populations, model accuracy for peak mosquito abundance can be improved by adding water station levels as a proxy for water habitat availability. Because they are a function of absorption and runoff processes on the ground, water levels measured at hydrological stations may be more reliable representations of standing water availability than precipitation measurements, which are known to be highly spatially variable [45]. Over-bank flow, as measured by river water gauges, is a direct quantification of flooding. Combining flood level information with temperature and daylight hours drivers better predicted the abnormally high mosquito population years than a predictive model that incorporated just the latter two variables.

We observed that the variations in the year-to-year abundance of mosquito populations are more accurately predicted when all three variables are considered, with better performance for stage gauge levels versus precipitation. The water station levels also appear to incorporate better the lag between mosquito populations and aquatic habitat availability. This may be because precipitation measures are taken during rainfall. In contrast, the water station levels are taken at locations where rainwater flows and collects in the following hours and days. Thus water stage gauge data could better inform standing water needed for *Culex* aquatic life stages.

Environmental data streams have been identified as important through both mechanistic and statistical studies. A comparison of Figure 6 and Figure 7 indicates that a data fusion process for which the environmental variables are used to inform the mechanistic dynamics of mosquito development yields a more accurate prediction. Our results also highlight water gauge data as an additional data source that can better capture year-to-year differences in mosquito abundance, particularly peak average mosquito abundance for years with flooding or extreme weather events.

Statistical models have elucidated which environmental factors are correlated to mosquito abundance. These studies have quantified the most important indicators and potential prediction accuracy. Although the statistical analysis can identify the significance of environmental factors related to mosquito abundance, they cannot show how the factors influence abundance biologically. Also, they cannot incorporate mitigation approaches and “what-if” scenarios to mosquito population control. Past studies have indicated that the nonlinear interactions between environmental variables and the mosquito life cycle are an essential part of the life-cycle and need to be included in the model. Mechanistic models can explicitly account for the interactions among the environmental parameters and mosquito abundance.

Using the process-based algorithmic approach as opposed to strictly applying a differential equation or statistical model allows for the fusion of environmental data feeds into the dynamics of mosquito development, thus capturing the seasonality of mosquito presence and the magnitude of the population during a flooding year. Thus we have a model that can analyze the effects of field-collected weather data on mosquito dynamics and test the impact of potential mitigation efforts. Somewhat unusual for many biological systems, our process-based model performs better than a standard statistical model in predicting mosquito abundance. This is likely because there are nonlinear and lagged interactions between mosquito population dynamics and the exogenous variables we use to predict them that are difficult for statistical models to capture.

Our mechanistic model’s ability to examine “what if” scenarios is also valuable for informing potential mitigation strategies and could be particularly useful when applied to the effects of climate change. Since the availability of standing water in cities is directly impacted by precipitation, it is essential for local agencies to be able to anticipate increases in precipitation and flooding, along with temperature change, to minimize the risk of WNV in the future effectively. Climate change has already been shown to cause more extreme weather patterns, which could lead to increased rainfall, flooding, and soil moisture in temperate regions like Toronto [46, 47]. Knowing the fluctuation in the size of mosquito populations during peak seasons of activity will establish a basis for the severity of public risk for contraction of the West Nile Virus.

This study is unique in fusing lab and field data with mosquito dynamics while incorporating competition in the larval stage and enabling predictions for all years continuously for which environmental data feeds are available. Unlike many comparable models, our model does not fit new initial conditions for each mosquito season; it runs year-round and uses diapaused mosquitoes from the previous season to start the next season simulation. We ran the model for 13 years across the entire Greater Toronto Area, instead of being restricted to a single year at a time. This is a more generalized approach to get a bigger picture of the mosquito population while depending on fewer fitted parameters.

Water levels within an urban area will almost certainly depend not only on weather but on water management strategies. Municipalities often have complex systems for managing stormwater as well as infrastructure for modulating water levels in municipal rivers and waterways [48, 49, 50, 51]. Stormwater management impacts the availability of breeding habitats for mosquitoes in urban settings and can impact the flushing of mosquito populations [52, 53]. The Greater Toronto Area focuses its water management strategy around local water management, working to divert water within regional sub-basins [54, 55, 52]. Previous stormwater management reports have noted that this strategy can cause localized flooding events [54]. 2013 was an extreme flood year in Toronto. Based on our results, the process-based model does better than the linear model of forecasting mosquito responses to this “extreme” event. In future work, it will be important to consider other areas with varying water management strategies to see if the importance of water station data holds.

Our process-based model is limited in that we do not have the mathematical theory to rigorously identify equilibria or quantify uncertainty outside of parameter sensitivity as we would with a differential equations model. The focus of this model, however, was not a mathematical analysis but a novel method for fusing environmental data along with a laboratory-based understanding of the progression of *Culex pipiens/restuans* life stages for creating a model which could replicate field observations of mosquito abundance. In particular, we wanted to capture better the year-to-year variation in abundance resulting from flooding or other environmental drivers. Our model has demonstrated that data alone is not as informative as the fusion of data and developmental dynamics. We have also highlighted that the type of data streams used matters. The use of water stage gauge measurements resulted in a more accurate prediction of the magnitude of population size throughout the years, particularly in flood years. An important next step will be testing the model at additional locations across North America.

## 5 Acknowledgements

Support for DS, DG, CM, CX, and JC provided by LDRD grants at Los Alamos National Laboratory. Support for JMH was from NSF Award #1563531. We thank the 36 local public health units and the service providers that trapped, counted, and recorded mosquitoes in the GTA and insightful conversations with them. Research presented in this article was supported by the Laboratory Directed Research and Development program of Los Alamos National Laboratory under project number 20190581ECR. The U.S. Department of Energy supported this work through the Los Alamos National Laboratory. Los Alamos National Laboratory is operated by Triad National Security, LLC, for the National Nuclear Security Administration of U.S. Department of Energy (Contract No. 89233218CNA000001).

## Notes

### Competing Interest Statement

The authors have declared no competing interest.

